# Human transferrin and lactoferrin cooperatively support *Neisseria meningitidis* colonization in the murine nasopharynx

**DOI:** 10.1101/2024.09.07.611816

**Authors:** Isaac S. Lee, Jamie E. Fegan, Elissa G. Currie, Anna Bojagora, Nelly Y. Leung, Derrick Rancourt, Muhamed-Kheir Taha, Anthony B. Schryvers, Scott D. Gray-Owen

## Abstract

*Neisseria meningitidis* is a human-restricted bacteria that is a normal nasopharyngeal resident, yet it can also disseminate, causing invasive meningococcal disease. Meningococci are highly adapted to life in humans, with human-specific virulence factors contributing to bacterial adhesion, nutrient acquisition and immune evasion. While these factors have been explored in isolation, their relative contribution during infection has not been considered due to their absence in small animal models and their expression by different human cell types not readily combined in either *in vitro* or *ex vivo* systems. Herein, we show that transgenic expression of the iron-binding glycoproteins human transferrin and lactoferrin can each facilitate *N. meningitidis* replication in mouse serum but that transferrin was required to support infection-induced sepsis. While these host proteins are insufficient to allow nasopharyngeal colonization alone, mice co-expressing these and human CEACAM1 support robust colonization. In this case, meningococcal colonization elicits an acute elevation in both transferrin and lactoferrin levels within the upper respiratory mucosa, with transferrin levels remaining elevated while lactoferrin returns to basal levels after establishment of infection. Competitive infection of triple transgenic animals with transferrin- and lactoferrin-binding protein mutants selects for bacteria expressing the transferrin receptor, implicating the critical contribution of transferrin-based iron acquisition to support colonization. These transgenic animals have thus allowed us to disentangle the relative contribution of three virulence factors during colonization and invasive disease, and provides a novel *in vivo* model that can support extended meningococcal colonization, opening a new avenue to explore the meningococcal lifestyle within its primary niche.

## INTRODUCTION

*Neisseria meningitidis* (*Nme*) is the etiological agent of invasive meningococcal disease, a rapidly progressing disease with severe clinical symptoms resulting in high fatality rates and frequent development of sequelae among survivors (1–4). Despite the devastating consequences of its invasive infections, *Nme* is a common resident of the human nasopharynx in healthy individuals, with an overall population prevalence of ∼10% and the duration of colonization often persisting for several months (4–8). *Nme* is human restricted and has become exquisitely adapted to life in humans. Indeed, many virulence factors that allow host cellular attachment or overcome host defense mechanisms strictly interact with the human form of their targets (9), which precludes their contribution to infection from being appreciated in non-human infection models.

During colonization, primary attachment to the apical surface of human mucosal epithelia is mediated by its bacterial type IV pilus, which extend and then retract to penetrate the mucus and overcome mucociliary clearance (10, 11). Then, an intimate secondary attachment is mediated by the colony opacity-associated (Opa) protein adhesins, a family of integral outer membrane proteins that are phase-variably expressed. Neisserial Opa proteins bind the human carcinoembryonic antigen-related cell adhesion molecules (hCEACAMs) on a variety of cell types (10, 12–15). Depending upon their expression level, which is further amplified by infection and/or cytokine exposure, CEACAM binding can promote cellular attachment, engulfment and/or transcellular transcytosis across polarized epithelia (12, 14, 16, 17). The influence of this interaction is apparent from the fact that transgenic mice that express hCEACAM1 become colonized by Opa-expressing *Nme* administered intranasally, an outcome not seen in wild type animals (18).

Once attached to the mucosal surface, bacteria must acquire essential nutrients such as carbon sources and transition metals to replicate and persist. However, some essential micronutrients are tightly regulated within the mammalian host to restrict microbial access and suppress their growth, a function known as nutritional immunity (19). In mammalian tissues, free iron is rapidly scavenged by the iron-binding glycoproteins transferrin (Tf) and lactoferrin (Lf), effectively creating an environment void of accessible quantities of this essential mineral. The pathogenic *Neisseria* overcome this restriction by expressing elegant receptor systems that directly bind to human Tf (hTf) and human Lf (hLf) to access their iron. These receptors are bi- partite transport systems, with a lipid-anchored hTf- or hLf-specific binding protein (TbpB or LbpB) on the surface of the outer membrane that captures iron-loaded versions of their respective targets and then couples it with an integral membrane β-barrel protein (TbpA or LbpA) that forms a gated channel to transport iron into the cell (19–22). The requirement for a utilizable iron source to *Nme* infection is well established as ferric dextran or hTf are routinely administered to animals prior to intraperitoneal injection of the bacteria for experimental sepsis studies (23–25) and transgenic mice that express hTf are highly susceptible to invasive meningococcal disease (26–28). For nasal colonization, which is necessarily the first stage of *Nme* infection, the relative contribution of endogenous hTf and hLf remains unclear, particularly when considering that the meningococci also have transport systems for host-derived heme (29) and siderophores from other bacteria (30). However, previous work has shown that intraperitoneal injection of hTf allowed *Nme* nasal colonization in very young (4-6 day) mice (31) and the administration of a very large bolus of hTf intraperitoneally followed by daily intranasal administrations supported *Nme* nasal and lung infection in adult mice (32).

In this study, we sought to dissect the relative contribution of physiologically relevant levels of hLf and hTf to meningococcal nasal colonization and sepsis through the establishment of mouse lines that endogenously express human CEACAM1, transferrin and lactoferrin transgenes. We observed that co-expression of these three factors supports prolonged meningococcal colonization that ultimately leads to the development of an *Nme-*specific humoral immune response. Further, we establish that hTf and hLf can each support *in vitro* growth in serum from transgenic mice and that hTf expression allowed prolonged bacterial persistence in the blood and more vigorous disease progression in systemically infected mice. When applying a bacterial genetic approach, we found that competitive intranasal infections with *Nme* lacking either component of the transferrin or lactoferrin receptors led to uniform selection for the strains that can use hTf, consistent with this becoming persistently elevated in mucosal secretions of infected animals. Thus, hLf appears to facilitate early colonization during a period of inflammation while hTf supports both mucosal infection and invasive meningococcal disease.

## METHODS

### Bacterial growth conditions

Wild type and mutant *N. meningitidis* serogroup B strain B16B6 (Table 1) were grown overnight on GC agar (Becton Dickinson (BD) #228930) supplemented with IsoVitalex (BD #B11876), followed by a subculture on GC agar (BD #228930) supplemented with IsoVitalex (BD #B11876) and 100 µg/mL of deferoxamine (Sigma-Aldrich #D9533-1G) for 4 h prior to infection for iron-starvation. GC agar (BD #228930) supplemented with IsoVitalex (BD #B11876) and VCNT inhibitor (BD #B12408) was used when recovering bacteria from mouse infection studies. All plates were incubated at 37°C with 5% CO2 for growth.

**TABLE 1:** *Neisseria meningitidis* strains and mutants

| Strain | Genotype<sup>a</sup> | Phenotype | Source |
| --- | --- | --- | --- |
| B16B6 | Wild type |  |  |
| B16B6 Δ*tbpA* | *tbpA::erm* | No TbpA expression | (20) |
| B16B6 Δ*lbpA* | *lbpA::cat* | No LbpA expression | (22) |
| B16B6 Δ*tbpB* | *tbpB::kan* | No TbpB expression | (20) |
| B16B6 Δ*lbpB* | *lbpB::kan* | No LbpB expression | (22) |

<sup>a</sup>*erm*, erythromycin resistance; *cat*, chloramphenicol resistance; *kan*, kanamycin resistance

### Mouse infection studies

All animal experimental procedures in this study were approved by the Local Animal Care Committee (LACC) at the University of Toronto (Protocol #20011319), in accordance with ethical and legal requirements under Ontario’s Animals for Research Act and the federal Canadian Council on Animal Care (CCAC). Mice were maintained in specific pathogen free conditions and were provided water and rodent chow *ad libitum*.

Human transferrin-expressing FvB mice (FvB hTf^+^) were generated by repeated backcrossing from the original C57BL6/SJLJ transgenic mouse line (33). Human lactoferrin expressing mice (hLf+) were generated for this study by the direct replacement of mouse Lf coding sequence with the human counterpart, subcloned from BAC clone RP2336G2, into G4 ES cells from an 129Sv x C57BL/6 cross. The resulting hLf gene is expressed from the mouse Lf promoter. These were repeatedly backcrossed for 12 generations onto the FvB background. hTf+ and hLf+ mice were crossed together to yield hTf^+^ hLf^+^ mice. These were further crossed with mice from the hCEACAM1^+^ line, which have been previously shown to support *Nme* colonization (18), to yield the various genotypes used in this study (Table 2). All mice were bred inhouse and littermate controls were used in all studies. Only males were used for sepsis studies, while both sexes were used for colonization studies.

**TABLE 2:** Mouse lines

| Strain | Expression | Description |
| --- | --- | --- |
| FvB hCCM1 | Human CEACAM1 | Transgenic expression of human CEACAM1 from human CEACAM1 promoter (57) |
| FvB hTf | Human transferrin | Transgenic expression of human transferrin from human transferrin promoter (33) |
| FvB hLf | Human lactoferrin | Genetic replacement of mouse lactoferrin gene with coding sequence from human lactoferrin, expressed from the endogenous mouse lactoferrin promoter (this study) |

**TABLE 3:**
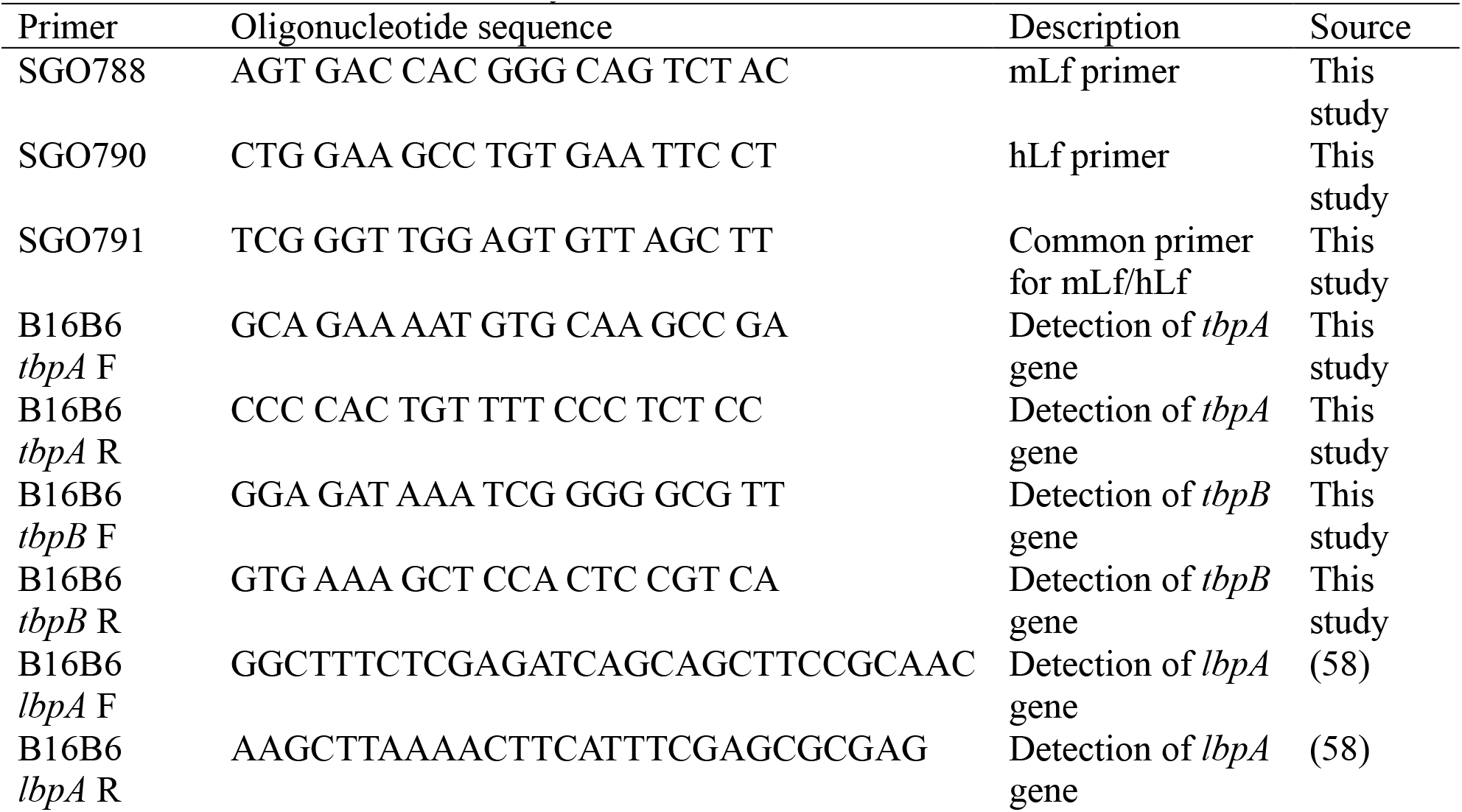
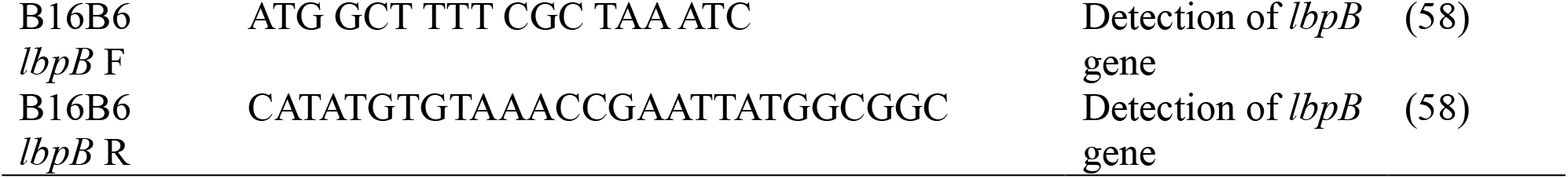
Primers used for the study

Nasal and systemic infections were performed as per previously described protocols (18). For nasal infections, isoflurane anaesthetized mice were intranasally infected with 1 x 10^8^ colony forming units (CFUs)/mL of *N. meningitidis* suspended in 10 μL phosphate buffered saline with 0.1 mg/mL Ca^2+^ and 0.2 mg/mL Mg^2+^ (PBS/Mg/Ca; Wisent). Mice were humanely euthanized at experimental endpoints of 1, 7 or 10 days post infection (pi), and cardiac bleeds, nasal lavages, and nasal swabs were collected. Nasal swabs were plated on GC agar supplemented with IsoVitalex (BD #B11876) and VCNT inhibitor (BD #B12408) for bacterial enumeration. Serum and nasal lavage samples were stored at -20 °C until analysis.

For invasive infections, mice were intraperitoneally infected with 1x10^8^ CFUs/mL of *N. meningitidis* suspended in 200 μL PBS/Mg/Ca. At indicated time points, 2.5 μL of blood was collected from the tail vein. The collected blood was 10-fold diluted in PBS/Mg/Ca and plated on GC agar supplemented with IsoVitalex and VCNT inhibitors for bacterial enumeration. Mice were monitored at 6-, 12-, 18-, 22-, 38- and 46-hours pi for clinical symptoms, at which point any mice that had reached clinical endpoint were humanely euthanized.

### Competitive infection

*N. meningitidis* TbpA^-^ and LbpA^-^ strains or TbpB^-^ and LbpB^-^ strains were grown overnight on plates as described above, followed by a 4 h iron-starvation subculture on GC agar supplemented with 100 µg/mL deferoxamine. The two strains were suspended in PBS/Mg/Ca, normalized to an equal OD600, and then plated for colony forming unit (CFU) enumeration. The bacterial suspensions and were mixed at 1:1 ratio according to their OD600 values, and 10 µL of the mixed inoculum containing ∼10^6^ CFU was used to intranasally inoculate mice. Mice were humanely euthanized at 3 days post-infection (PI), and nasal swabs were plated on GC agar supplemented with IsoVitalex (BD #B11876) and VCNT inhibitor (BD #B12408) for bacterial enumeration. Bacterial DNA was extracted from each CFU recovered using the QuickExtract DNA extraction solution (Epicentre, # QE09050) and each colony was screened for the presence of *tbpA* and *lbpA* genes, or *tbpB* and *lbpB* genes via PCR. Competition index was calculated using the following equation, which corrects for the possible concentration differences between optical density estimates and plated inoculum concentrations:

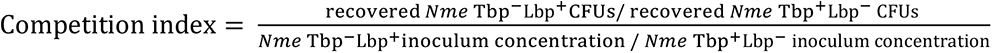

### In vitro growth curves

*N. meningitidis* was prepared overnight on plates, followed by the 4 h iron-starvation subculture step described above. Bacteria were swabbed from plates and resuspended in PBS/Mg/Ca. Growth curves were performed in U-bottom 96-well plates (Greiner Bio-One #650180), with each well containing 65 μL of RPMI (Wisent #3500-000-CL), 25 μL of transgenic mouse serum or human serum (Sigma-Aldrich, H4522-100ML) and 15 μL of bacterial suspension to obtain a starting OD600 of 0.1, and plates were sealed with Breathe-Easy plate seal (Sigma-Aldrich #Z380059-1PAK). Bacterial growth was measured every 30 min for 18 h using a Cytation 5 cell imaging multimode reader (BioTek).

### ELISAs for relative abundance of hLF

Terminal serum, nasal lavage fluids and lung lavage fluids were collected from mice as described previously (18). Mouse neutrophils were isolated from bone-marrow, stimulated with PMA to induce degranulation, and supernatants were collected for ELISAs as previously described (34, 35). Nunc MaxiSorp 386-well ELISA plate (Thermofisher, 12-565-347) was coated with neat samples. Rabbit α-human lactoferrin polyclonal antibody (Sigma-Aldrich, L3262) was used for detection, with the plate washed and subsequently probed using 1:5000 of goat α-rabbit IgG – HRP antibody (Jackson Immunoresearch, 111-035-144).

### ELISAs

Terminal serum and nasal lavage fluids were collected from mice as described previously (18) and stored at -20 °C for analysis. Samples were diluted 1:20 for the detection of mouse IgG (Biolegend, 432401), mouse myeloperoxidase (MPO; R&D Systems, DY3667) and human lactoferrin (Abcam, ab200015), and diluted 1:500 for the detection of human transferrin (Abcam, ab187391).

### Whole Cell ELISAs

*Nme* was grown overnight as described above, heat inactivated at 65 C° for 1 hour, and subsequently added to 386-well plates (20 μL/well) at the optical density (OD600) of 0.2 before drying overnight at room temperature (RT) in a biological safety cabinet. The effectiveness of heat inactivation was confirmed via plating the treated suspension on GC Agar + IsoVitalex. The coated plate was blocked with 5% bovine serum albumin (BioShop, ALB001) at RT for 1 h.

Mouse serum from infected mice were added to the plates (1:50 dilution) and incubated for 2 h at RT. Plates were washed and subsequently probed using 1:5000 and 1:10000 dilutions of α-mouse IgA-AP (Abcam, ab97232) and α-mouse IgG-HRP antibodies (Jackson ImmunoResearch, 115- 035-003), respectively.

### In vivo depletion of neutrophils

Neutrophils were depleted from mice by intraperitoneal injection of 200 µL 1.25 mg/mL anti- GR1 monoclonal antibody (purified from RB6-8C5 hybridoma, (18)) at 72 and 24 hours prior to infection. Neutrophil depletion was confirmed by myeloperoxidase (MPO) ELISA.

## RESULTS

### Functional replacement of mouse lactoferrin gene with that encoding human lactoferrin

While hLf is considered a major source of iron for *Nme* during nasopharyngeal colonization, *in vivo* studies to consider the physiological relevance of hLf have not been possible due to the strict specificity of Lbps for human-derived Lf. Thus, we generated a transgenic mouse line that replaces the mouse lactoferrin gene with that of human (Fig. 1A). The transgenic hLf mice effectively produce and release hLf within the respiratory mucosa as expected. Only trace amounts of hLf are detected in sera, consistent with it being thought to be only released upon systemic inflammation due to neutrophil degranulation (Fig. 1B). Neutrophil expression in neutrophil granules was evident by the fact that supernatants from PMA-stimulated bone marrow-derived neutrophils contained increased hLF relative to the purified but unstimulated cells (Fig 1C).

**Figure 1.**
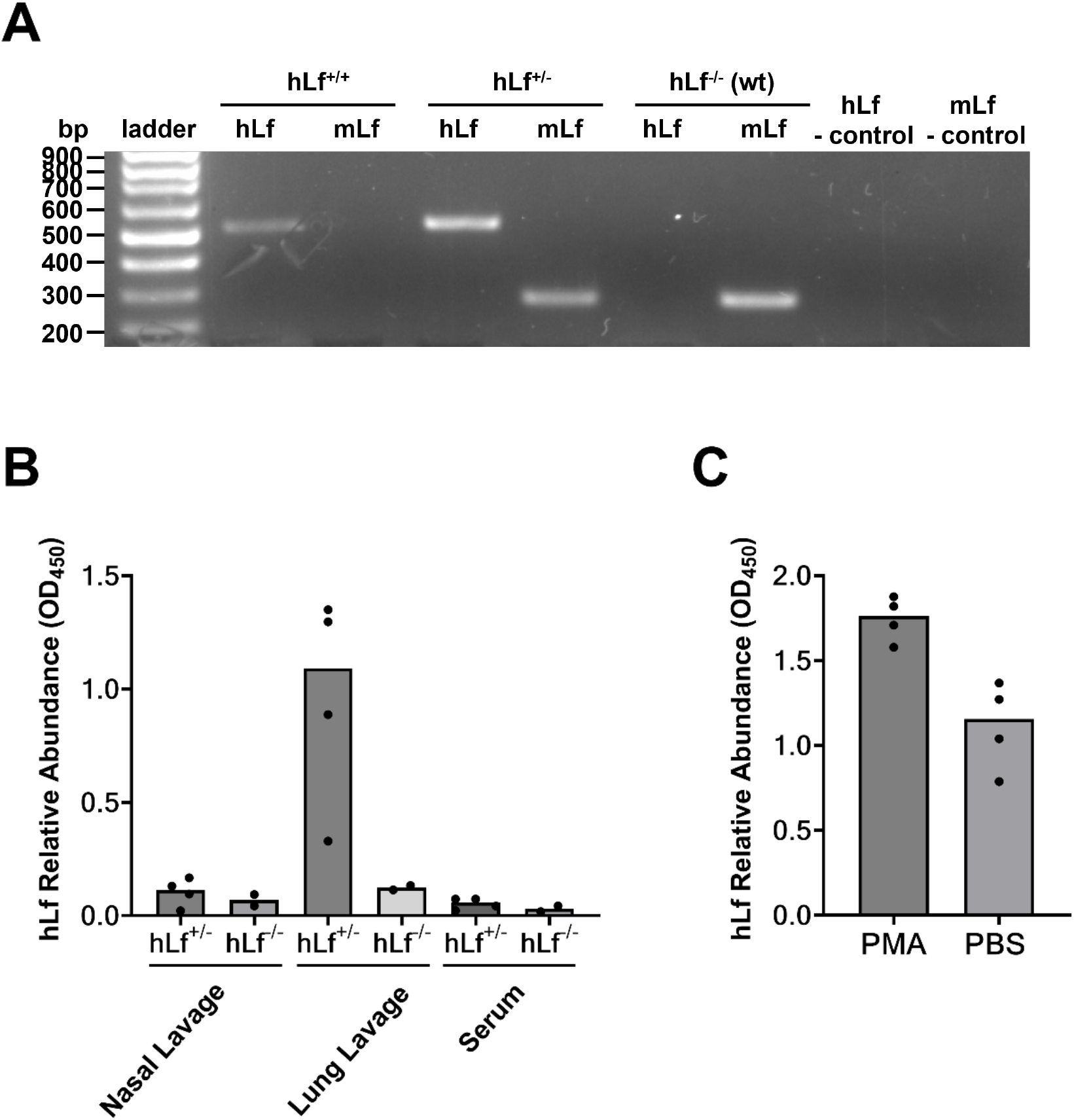
Validation of human lactoferrin expression in transgenic mice. (A) PCR amplified bands of hLf and mLf gene fragments for hLf^+/+^, hLf^+/-^, and hLf^-/-^ mice. (B) Relative hLf levels in nasal lavage, lung lavage and serum of heterozygous transgenic hLf or wild type mice were evaluated by hLf ELISAs. Data is from 4 mice for each genotype. (C) Neutrophil degranulation releases hLf. Bone-marrow derived neutrophils of hLf heterozygous transgenic mice were stimulated with PMA or mock-treated with PBS to induce degranulation. The supernatants were run on human lactoferrin-specific ELISAs to evaluate the relative abundance of hLF post-degranulation. Data are from 2 mice with technical replicates. Abbreviations: hLf, human lactoferrin; PMA, phorbol 12-myristate-13-acetate; PBS, phosphate buffered saline; OD, optical density. Bars represent median.

### Sera from human transferrin or lactoferrin expressing mice is sufficient to support meningococcal growth

hTf expressing mice have been shown to sustain *Nme* systemic infection (36–38), while such capability has not been previously determined for the newly generated hLf mice. To evaluate the relative ability for endogenously expressed hTf and hLf to support meningococcal growth, liquid cultures were established with serum from transgenic mice or wild type (WT) littermate controls as the sole source of iron. *Nme* was able to grow with human serum as a sole iron source, while little growth was apparent with WT mouse serum (Fig. 2A). Growth in hTf^+/-^ mouse serum was comparable to that of human serum, consistent with previous literature (37). *Nme* was also able to grow with hLf^+/-^ mouse serum as the sole iron-source (Fig. 2A). This indicates that the human-derived proteins are functionally expressed and available to the meningococci.

**Figure 2.**
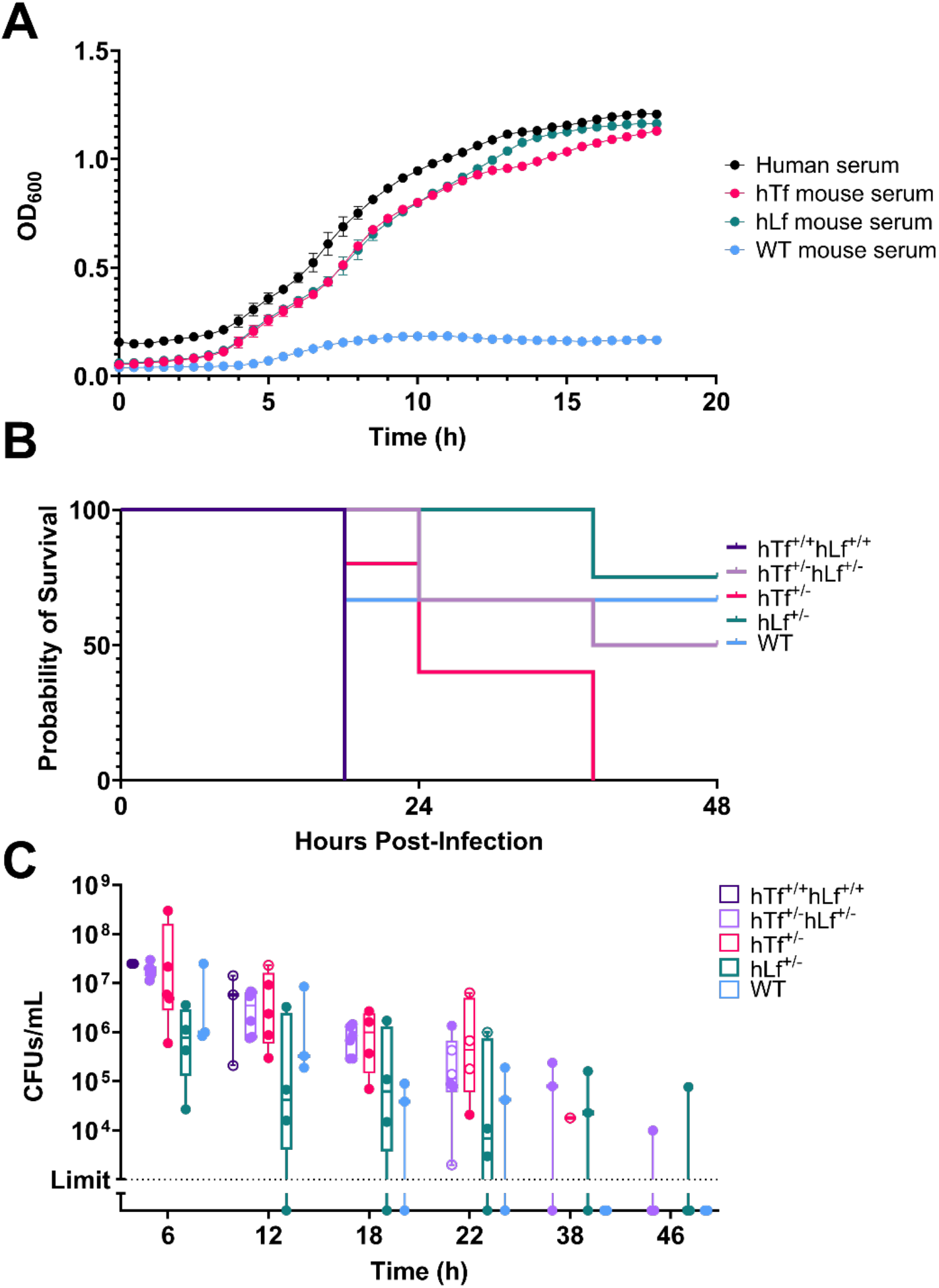
Ability of hTf and hLf transgenic mouse serum to support *N. meningitidis* growth. (A) Human serum or that from hTf^+/-^, hLf^+/-^ or wild type mice (20% v/v) were used as sole iron source to facilitate *N. meningitidis in vitro* growth in RPMI media. (B and C) hTf^+/+^ hLf^+/+^, hTf^+/-^ hLf^+/-^, hTf^+/-^, hLf^+/-^, or wild etype mice were intraperitoneally infected with 5x10^8^ CFU/mouse of *N. meningitidis*. (B) Survival and (C) Bacterial burden in infected mice. Burden is represented by colony forming units (CFUs) recovered per mL of blood collected via tail vein sampling, collected at 6-, 12-, 18-, 22-, 38- and 46-hours post-infection for surviving mice only. Clinical scores were assessed based on pre-determined criteria, with a score of 0 representing no symptoms to 10 representing clinical endpoint. Open circles represent mice at endpoint, after which there is no value plotted for that animal. Abbreviations: WT, wild type; hTf, human transferrin; hLf, human lactoferrin; CFUs, colony forming units. (A) Data are means +/- SD and are representative of 3 independent experiments. (B and C) 3-6 mice per group.

*Nme* is not typically capable of replicating or causing symptomatic bloodstream infection in mice without the addition of an accessible iron source such as iron-dextran or hTf (23–25). Systemic meningococcal infection was performed in transgenic mice expressing hTf, hLf or both to determine whether these iron sources were sufficient to support bacterial persistence and disease progression (Fig. 2B-C). The highest lethality was seen in mice expressing hTf, with all hTf^+/+^ hLf^+/+^ mice reaching clinical endpoint by 18 hours pi and all hTf^+/-^ hLf^-/-^ mice reaching clinical endpoint by 38 hours pi. The survival of mice that are heterozygous for hLf but lacking hTf (75%) was not different than the wild type animals (67%), implying that hLf does not effectively support meningococcal disease progression. Mice that are heterozygous for both hTf and hLf also, unexpectedly, had similar clinical outcomes to the wild type animals (50%). Notable in this regard, the bacterial burden in blood remained higher for longer in all transgenic versus wild type animals. The bacteremia was found to persist for extended durations in the transgenic animals, being still detectable in some hTf^+/-^, hLf^+/-^ and hTf^+/-^hLf^-/-^ at 38 and 46 h pi (homozygous hTf^+/+^hLf^+/+^ had all reached endpoint by this point), while WT mice had no recoverable *Nme* after 22 h pi. However, the mice that succumbed did not have higher bacterial CFUs recovered from blood at the time point prior to their reaching the clinical endpoint, suggesting that the same burden of meningococci is more virulent in the presence of an iron source and/or that the bacteria may also be replicating in a niche other than blood.

### Expression of human transferrin and lactoferrin extend the duration of nasopharyngeal colonization by *N. meningitidis*

Despite hCEACAM1 expression in mice being sufficient to allow short-term (<7 day) nasal colonization by *Nme* (18), this model is unable to support longer term persistence of the bacteria, limiting the breadth of study that is feasible (5, 6, 18). We hypothesized that the limited duration was due to the inability of the bacteria to acquire necessary micronutrients including iron. To directly assess whether hTf and/or hLf could sustain prolonged infection, we crossed the three mouse lines to generate transgenic animals expressing varied combinations of hCEACAM1, hTf and/or hLf.

With the exception of 2 hTf-expressing mice with low bacterial recovery, mice lacking hCEACAM1 were not colonized by *Nme* (Fig. 3A). Consistent with previous work (18), *Nme* was not recovered from hCEACAM1 mice at day 7 post infection. However, the co-expression of hTf and/or hLf with hCEACAM1 proved effective in extending the duration of meningococcal colonization of the murine nasopharynx beyond 7 days, with over half of the animals expressing both hLf and hTf remaining infected at 10 days pi. While the proportion of mice that remained colonized decreased from 7 to 10 days, it is that the burden of *Nme* in animals that remain infected is similar at these two time points (Fig. 3A). At each time point, the proportion of animals that express both hTf and hLf was always higher than those that express either protein alone, suggesting that they both contribute to bacterial persistence. Further, animals expressing hCEACAM1 with hTf contained significantly greater levels of α-*Nme* IgG compared to animals expressing hCEACAM1 alone, suggesting that augmented persistence may lead to stronger adaptive response in this model (Fig. 3B).

**Figure 3.**
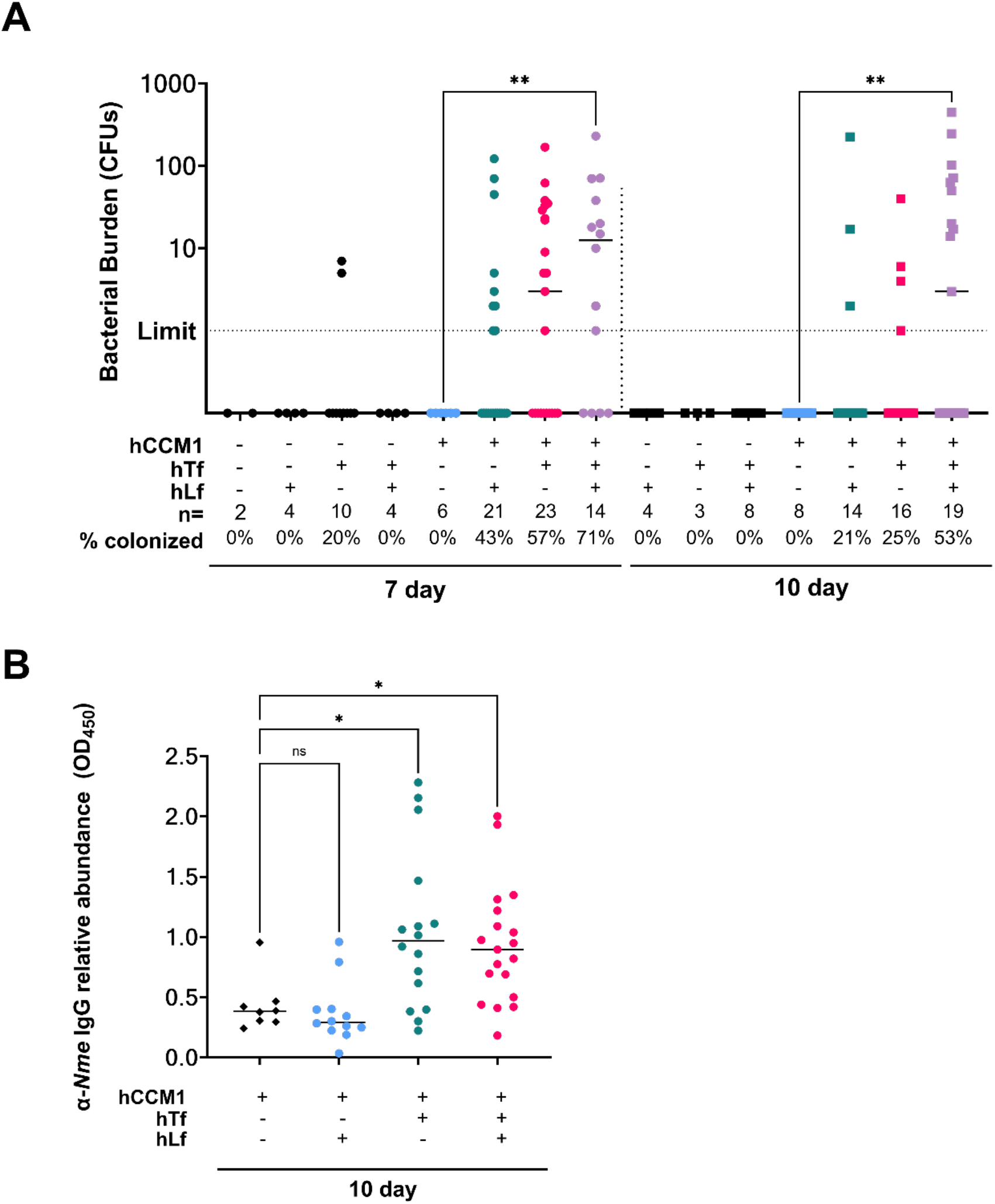
hTf and hLf promote sustained *N. meningitidis* nasopharyngeal colonization in mice. Indicated mouse lines were nasally infected with 10^6^ CFU/mouse of *N. meningitidis*. Bacterial enumeration was done 7- and 10-days post-infection via nasal swabs. (A) Colonization rates and CFUs for 7- or 10-days post-infection per genotype. Data are pooled from 4 independent experiments. (B) *Nme-*specific IgG present in serum at 10 days post-infection; bars represent group median. Kruskal-Wallis test with multiple comparisons. *p < 0.05, **p< 0.01,. hCCM1, human CEACAM1; hTF, human transferrin; hLF, human lactoferrin.

Utilization of human transferrin is more advantageous than human lactoferrin for *N. meningitidis* colonization during murine nasopharyngeal colonization To confirm that the effect of hTf and hLf expression was due to their ability to act as an iron source for the meningococci, we next explored how the specific absence of individual receptor proteins influenced *Nme* infection. Given that the integral membrane proteins, TbpA and LbpA, are essential for neisserial iron acquisition from their respective host proteins while the lipid-anchored TbpB and LbpB are not essential but do increase the efficiency of this process (22, 39, 40), we validated the phenotype of mutant strains lacking each protein individually by comparing their ability for each to grow in media containing either hTf or hLf as the sole iron-source. As expected, Δ*tbpA* and Δ*lbpA* strains were unable to grow in hTf or hLf containing media, respectively, but were able to utilize the alternate iron source (Fig. 4Ai, Aiii). In contrast, the Δ*tbpB* and Δ*lbpB* strains were able to grow with hTf and hLf respectively, though at a rate intermediate between that of the wild type and the Δ*tbpA*/Δ*lbpA* strains (Fig. 4 Aii, Aiv). These results are consistent with the previous *in vitro* studies.

**Figure 4.**
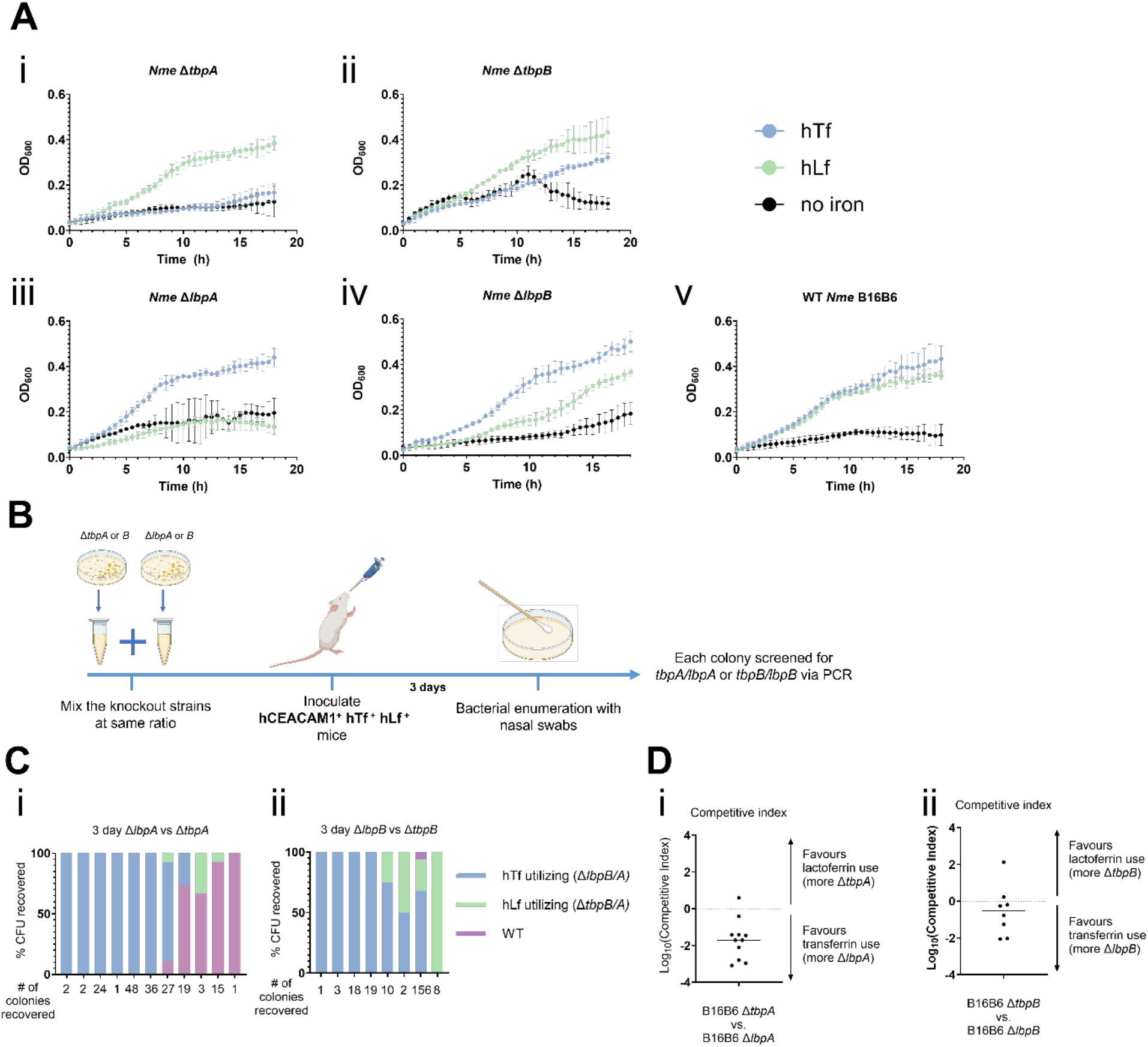
The relative importance of hTf and hLf during murine nasopharyngeal colonization of *N. meningitidis.* (A) *N. meningitidis* (i) *ΔtbpA*, (ii) *ΔtbpB*, (iii) *ΔlbpA*, (iv) *ΔlbpB* or (v) wild type strains were grown in RPMI with either 0.1 mg/mL hTf, 0.1 mg/mL hLf, or no iron. (B) Schematic overview for the competitive infection experiments. (C) *N. meningitidis* (i) *ΔtbpA* and *ΔlbpA* or (ii) *ΔtbpB* and *ΔlbpB* were used to inoculate hCEACAM1^+/-^ hTf^+/-^ hLf^+/-^ mice. Bacterial enumeration was done at 3 days post-infection via nasal swabs. Percentage of each mutant colonies recovered per mouse are shown. (D) Competitive index of (i) *ΔtbpA* and *ΔlbpA* competition infection at 3 days pi, or (ii) *ΔtbpB* and *ΔlbpB* competition infection at 3 days pi. Competitive index = (*ΔtbpA* or *Δtbp*B:*ΔlbpA* or *ΔlbpB*)/ (*ΔtbpA* or *ΔtbpB* inoculum concentration/*ΔtbpA* or *ΔtbpB* inoculum concentration). Log10[Competitive index] > 0 = selective advantage for *ΔtbpA* or *ΔtbpB* (hLf utilizers); Log10[Competitive index] < 0 = selective advantage for *ΔlbpA* or *ΔlbpB* (hTf utilizers). For the purpose of the calculation, value of 0.1 was used when no colonies were recovered for either mutant.

Having verified the phenotype of the mutant strains, a competition infection was performed using Δ*tbpA* and Δ*lbpA* strains to infect transgenic mice co-expressing hCEACAM1, hTf and hLf. Three days post infection, 6 of 11 mice were solely colonized by the hTf utilizing Δ*lbpA* strain, while a mixed population comprised of Δ*tbpA*, Δ*lbpA* and, unexpectedly, wild type *Nme* were recovered from 4 of 11 mice, and one mouse was solely colonized by WT *Nme* (Fig. 4Ci). The genotype of each colony was confirmed by PCR, so the recovered WT *Nme* are likely to be revertant mutants that had acquired a wild type *tbpA* or *lbpA* by horizontal gene transfer. To numerically evaluate the selective advantage, the competition index was calculated as the ratio of Δ*tbpA* to Δ*lbpA* colonies recovered and then normalized for the differences in the concentration of respective mutants present in the inoculum (as determined by plating). In this case, Log10[Competition index] value of 0 represents no selective advantage, a positive value represents an advantage with hLf utilization, and a negative value represents an advantage for hTf-expressing strains. This clearly revealed that transferrin utilizing *Nme* had a survival advantage during nasopharyngeal infection (Fig. 4Di).

To consider whether the reduced efficiency of iron acquisition in the absence of the lipoprotein component of the hTf or hLf receptors had a physiologically relevant effect, we performed a competition infection with the Δ*tbpB* and Δ*lbpB* strains. At 3-days pi, 4/8 mice were exclusively colonized by hTf-utilizing Δ*lbpB* strain, 3/8 mice were co-colonized by a combination of of Δ*lbpB* and Δ*tbpB* strains (Fig 4Ciii) and 1/8 mouse exclusively carried the Δ*tbpB* strain (Fig 4Ciii). Only a single colony of wild type *Nme* was recovered when these strains were combined. Collectively, these results suggest that transferrin utilization is crucial and imposes a stronger selective advantage than lactoferrin utilization during nasopharyngeal colonization by *Nme*.

### Transferrin and lactoferrin are released to the mucosal surface by the host in response to *Nme* challenge

It was unexpected that transferrin utilization would be strongly selected over lactoferrin utilization given that hLf^+/-^ hCEACAM1^+/-^ mice supported similar colonization by *Nme* at 7 days pi. We explored whether this advantage for hTf utilization is a function of differences in the abundance of transferrin and lactoferrin in the mucosa during *Nme* infection. For this, hTf and hLf levels were quantified by ELISA for nasal lavages recovered from hTf^+/-^hLf^+/-^hCEACAM1^+/-^ mice that had been intranasally administered either PBS/Mg/Ca or *Nme*. Both hLf and hTf levels are markedly higher 1 day after *Nme* challenge compared to the levels in mock infected (PBS/Mg/Ca treatment) animals, reaching approximately 0.2 and 1 μg/mL, respectively (Fig. 5Ai, iii). Notably, hLf levels returned to that seen in the PBS-treated animals by 7 and 10 days pi, while hTf levels remained elevated at 10 days pi.

**Figure 5.**
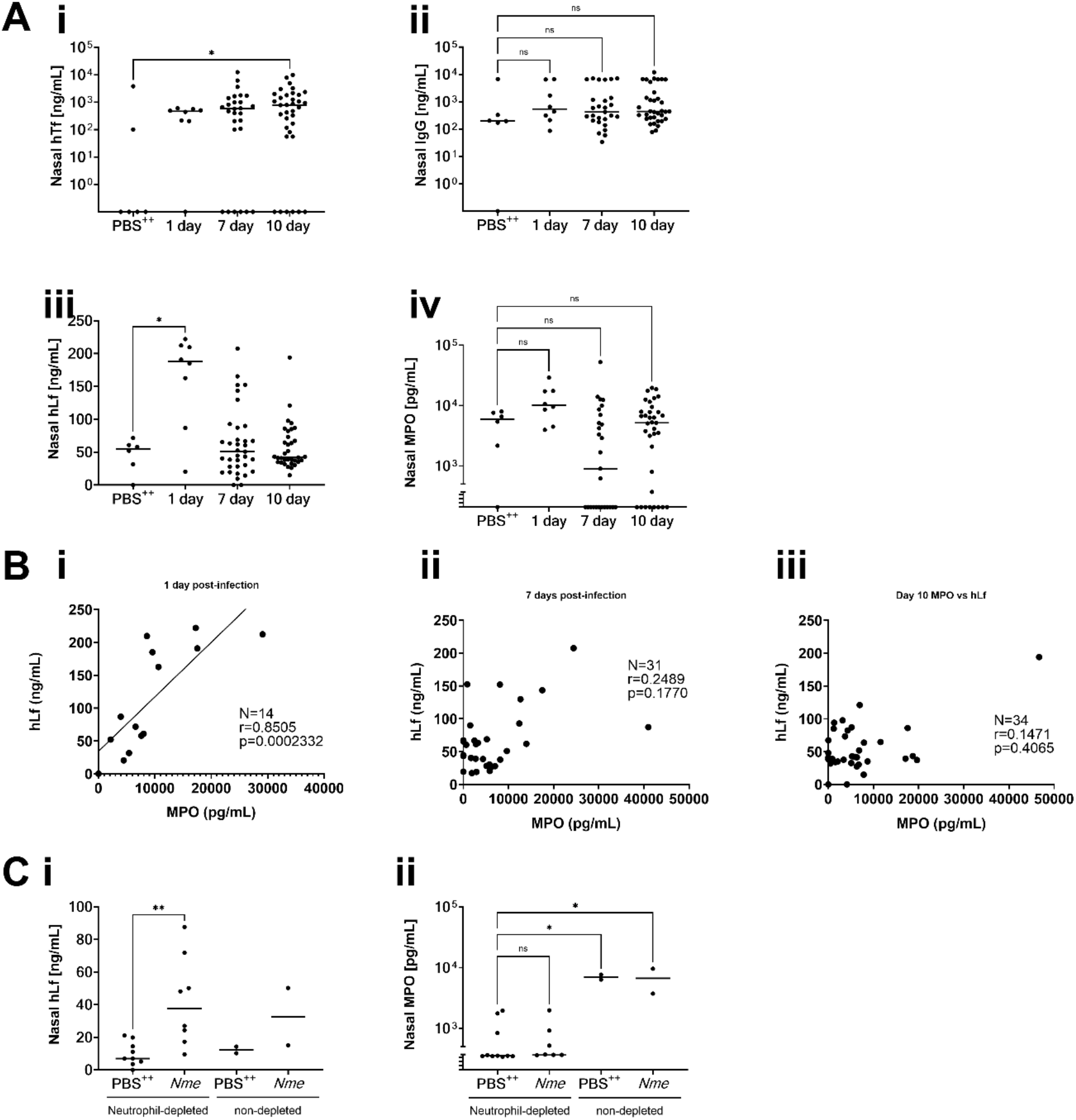
Transferrin and lactoferrin levels are elevated in the nasopharynx of hTf and/or hLf expressing mice upon infection. A) hCEACAM1^+/-^ hTf^+/-^ hLf^+/-^, hCEACAM1^+/-^ hTf^+/-^, or hCEACAM1^+/-^ hLf^+/-^ mice were inoculated with PBS^++^ or *N. meningitidis*, and nasal lavages were collected 1 day post-infection. In a separate experiment, hCEACAM1^+/-^ hTf^+/-^ hLf^+/-^, hCEACAM1^+/-^ hTf^+/-^, or hCEACAM1^+/-^ hLf^+/-^ mice were inoculated with *N. meningitidis* and nasal lavages were collected either 7- or 10-day post-infection. Nasal lavages were used to quantify myeloperoxidase (MPO), IgG, human lactoferrin and human transferrin levels by ELISA. B) Correlation between hLf and MPO in nasal lavages of *N. meningitidis* infected mice at 1-, 7- and 10-days pi. Spearman correlation. C) hCEACAM1^+/-^ hTf^+/-^ hLf^+/-^, hCEACAM1^+/-^ hTf^+/-^, or hCEACAM1^+/-^ hLf^+/-^ mice were depleted of neutrophils via IP injection of α-GR1 Ab at 72- and 24-hours prior to infection. Neutrophil depleted mice were inoculated with PBS^++^ or *N. meningitidis*, and nasal lavages were collected 1-day post-infection. Nasal lavages of the depletion experiment were run alongside samples from the previous non-depleted experiment to quantify myeloperoxidase and human lactoferrin levels by ELISA. Kruskal-Wallis test with multiple comparisons. *p < 0.05, **p< 0.01.

Transferrin is not typically considered a mucosal protein, as it originates from the liver and primarily traffics through the blood into the tissues (41). We considered that the rapid recruitment of hTf into the nasopharyngeal lumen may result from transudation from the bloodstream in response to inflammation, which can reduce the integrity of epithelial barrier function (42). To explore this, luminal IgG was measured in the lavages. A modest increase in nasal IgG levels was apparent in *Nme-*infected animals, though this difference was not statistically significant. While both active and passive processes can facilitate IgG passage across the epithelia, it is not affected by *Nme* infection in a manner reflecting that seen with hTf.

The rapid increase in hLf within the luminal washes could be due to local production of the protein within mucosal secretions or via neutrophil recruitment given that these innate phagocytes can robustly respond to infection and release lactoferrin from their secondary granules upon exposure to microbial stimuli (43–45). We first explored the possible link between hLf levels and the neutrophil response by measuring neutrophil-derived myeloperoxidase (MPO) levels in the nasal lavages. This revealed a significant correlation between MPO levels and hLf levels at 1 day pi, but no correlation at the later time points (Fig 5B). To determine whether this correlation is due to a causal relationship between neutrophil activity and hLf levels, we immunologically depleted neutrophils from the animals prior to meningococcal challenge, and collected nasal lavages at 1 day pi. While the MPO response no longer occurred in the neutrophil-depleted animals, hLf levels remained elevated after meningococcal challenge (Fig. 5C), suggesting that the increased hLf was instead a localized response by the epithelia.

## DISCUSSION

Investigation into meningococcal nasal carriage has historically been challenging due to the lack of small animal models that can adequately support stable colonization of this human-restricted bacteria. Herein, we generated an improved model of *N. meningitidis* nasopharyngeal colonization by combining transgenic expression of two human iron-binding glycoproteins, transferrin and lactoferrin, with the CEACAM1-humanized mouse model. While hCEACAM1 expression is essential for nasopharyngeal colonization by *Nme*, co-expression of hTf and hLf substantially increased the bacterial burden and persistence of infection beyond that seen with hCEACAM1 alone. Co-infection with meningococcal mutants that lack individual components of the transferrin or lactoferrin receptor reveal that there is a clear selective advantage for strains that can use hTf, though the burden of infection is still higher when animals express both iron sources. This bias is consistent with our observation that hTf becomes abundantly available in the infected mucosal tissues for the duration of infection, while hLf is only transiently increased after infection.

Prior work had already established the impact of the endogenous expression of the hTf transgene to support invasive meningococcal disease following intraperitoneal inoculation or nasally administered to mice that had been pre-infected with a sublethal dose of influenza virus (26–28). Our results reinforce the importance of hTf during invasive disease by showing that hLf expression does not have a similar effect. Our finding that hTf is key to support mucosal colonization is, perhaps, unexpected given the classical assumption that the bacterial transferrin receptor supports invasive disease while the lactoferrin receptor allows for the scavenging of iron on mucosal tissues, where lactoferrin can be produced by glandular epithelia. However, the essential requirement for access to hTf iron is intuitive given that all pathogenic *Neisseria* express the Tbps while almost half of *N. gonorrhoeae* lack Lbps (46), and that *Haemophilus influenzae* (47) and other host-restricted respiratory pathogens of the Pasteurellaceae (48) express transferrin but not lactoferrin receptors. The sustained increase in transferrin that we observed within the nasal lumen throughout the duration of infection helps to explain the utility of microbes targeting transferrin. It is satisfying that the differences apparent in these humanized mice reflects those seen in humans, with prior work reporting that hTf is ∼10-fold more abundant than hLf in bronchoalveolar fluids (49). It thus seems likely that Lbps facilitate growth during the transient inflammatory response elicited in the early stages of infection, when neutrophils are abundant. Consistent with this premise, LbpB also protects against the bactericidal activity of lactoferrin’s carboxyl-terminal sequence, which can be proteolytically released at sites of inflammation to form lactoferricin, a cationic antimicrobial peptide (50).

Neutrophil depletion prior to infection had no significant effect on the surge of hLf that was apparent in nasal washes 1 day post-infection, suggesting that this was locally-derived from the epithelia. This is also consistent with lactoferrin transcription being induced upon celluar exposure to lipopolysaccharide and regulated by the transcription factor nuclear factor kappa B (NF-κB), which governs the inflammatory response (51). Given that mucosal inflammation promotes reorganization of epithelial intercellular junctions, which disrupts the barrier integrity (42), it would be reasonable to suggest that transferrin accumulation within the nasopharyngeal lumen results from transudation into the nasopharyngeal lumen. However, a similar increase in IgG concentrations was not apparent, so other processes may also promote transferrin movement into the mucosa. For example, macrophages and neutrophils both have the potential to express transferrin (52, 53), though the transient nature of the inflammatory response that occurs during *Nme* colonization makes it questionable as to whether this could sustain hTf at the levels seen. In considering the host cellular response, neisserial infection has been shown to downregulate expression of the human transferrin receptor in epidermoid cells (54), making it feasible that transferrin passively accumulates by not being removed from the mucosal lumen. Also, provocatively, *Helicobacter pylori* has been observed to dysregulate the human transferrin receptor expressed by gastric epithelial cells, causing active transcytosis of hTf from the basolateral to apical surface (55). If *Nme* has a similar capacity, then this would lead to hTf accumulation in the mucosal surface as observed here.

Beyond insights regarding the relative contribution of hTf and hLf to infection of nasopharyngeal tissues, our findings establish the utility of these humanized mice for studying meningococcal biology and the host response to asymptomatic colonization, which naturally precedes invasive meningococcal disease. The frequent isolation of wild type colonies expressing both Tbp and Lbp receptors after co-infecting with mutants that were all defective in one or the other, a dramatic example of the effectiveness of natural competence of these bacteria (56), reflects the opportunity to study horizontal genetic transfer or other aspects of the lifestyle of these bacteria within their physiologically relevant mucosal niche.

## Acknowledgements

We thank Laura-lee Caruso for technical assistance and Cynthia Xinyi Guo for helpful discussions through the course of this study.

This research was supported by Canadian Institute for Health Research (CIHR) grant PJT- 153177 and National Institutes of Health grant R01-AI125421-01A1. S.D.G. is supported by the Canada Research Chair Program. I.S.L. received support from the CIHR Canadian Graduate Scholarship – Master’s Award Program and E.G.C. was supported by the CIHR Doctoral Award program.

## Notes

### Competing Interest Statement

The authors have declared no competing interest.

